# Environmental predictors of electroactive bacterioplankton in small boreal lakes

**DOI:** 10.1101/2022.03.26.485925

**Authors:** Charles N. Olmsted, Roger Ort, Patricia Q. Tran, Elizabeth A. McDaniel, Eric E. Roden, Daniel R. Bond, Shaomei He, Katherine D. McMahon

**Affiliations:** Department of Molecular and Environmental Toxicology, University of Wisconsin, Madison, 1550 Linden Drive, Madison WI, 53706; Trout Lake Station, Center for Limnology, University of Wisconsin-Madison, 10810 County Highway N, Boulder Junction, WI 54512, USA; Department of Bacteriology, University of Wisconsin, Madison, 1550 Linden Drive, Madison WI, 53706, USA; Department of Integrative Biology, University of Wisconsin - Madison, Madison, WI; Microbiology Doctoral Training Program, University of Wisconsin - Madison, Madison, WI; Department of Geoscience, University of Wisconsin, Madison, 1215 West Dayton Street Madison, WI 53715, USA; Department of Plant and Microbial Biology and BioTechnology Institute, University of Minnesota, 1479 Gortner Ave, St. Paul MN 55108; Department of Civil and Environmental Engineering, University of Wisconsin, Madison, 1550 Linden Drive, Madison, WI, 53706, USA

## Abstract

Extracellular electron transfer (EET) by electroactive bacteria in anoxic soils and sediments is an intensively researched subject, but EET’s function in planktonic ecology has been less considered. Following the discovery of an unexpectedly high prevalence of EET genes in a bog lake’s bacterioplankton, we hypothesized that the redox capacities of dissolved organic matter (DOM) enrich for electroactive bacteria by mediating redox chemistry. We developed the bioinformatics pipeline FEET (Find EET) to identify and summarize EET proteins from metagenomics data. We then applied FEET to several bog and thermokarst lakes and correlated EET protein occurrence values with environmental data to test our predictions. Our results provide evidence that DOM participates in EET by bacterioplankton. We found a similarly high prevalence of EET genes in most of these lakes, where oxidative EET strongly correlated with DOM. Numerous novel clusters of multiheme cytochromes that may enable EET were identified. Taxa previously not considered EET-capable were found to carry EET genes. We conclude that EET and DOM interactions are of major ecological importance to bacterioplankton in small boreal lakes, and that EET, particularly by methylotrophs and phototrophs, should be further studied and incorporated into both conceptual and quantitative methane emission models of melting permafrost.

## INTRODUCTION

Aquatic microbes influence Earth’s atmosphere directly and indirectly through fermentation, respiration, photosynthesis, and other metabolisms all requiring the transfer of electrons. Water provides a medium for integrating solutes involved in these processes, yielding high metabolic turnover at rates that make water bodies hubs for the landscape’s biogeochemical processing. We present evidence that in small freshwater boreal lakes planktonic metabolisms may often be more connected than previously realized—electroactively through extracellular electron transfer (EET). EET is the conduction of electrons through the outermost membranes of cells, allowing metabolisms like respiration to make use of solute or solid electron acceptors without having to transport them into the cell.

In and on structured environments such as rocks, soils, sediments, and conductive surfaces there is ample evidence to demonstrate that, ubiquitously, electroactive organisms can be found using EET to connect internal electrochemistry to the external environment (Coates *et al*., 2002; Roden *et al*., 2010; Tang *et al*., 2019; McAllister *et al*., 2020; Keffer *et al*., 2021; Rowe *et al*., 2021) or to other bacteria (Hegler *et al*., 2008; Shi *et al*., 2016; Beyenal *et al*., 2017; Ishii *et al*., 2018). This led studies to focus on electron transfer to solid-phase substances. However, the ability of bacteria to reduce soluble electroactive substances is well-known, and this raises the possibility of their use as substrates for EET rather than as mediators or electron shuttles to a solid substrate (Lovley *et al*., 1991; Bond and Lovley, 2005; Li *et al*., 2019). For example, a surprisingly high number of genes encoding multiheme cytochromes (MHCs) and other putative EET proteins were found in bacteria inhabiting the water column of Trout Bog Lake, a humic lake in WI, USA, and researchers pointed to the high electron accepting capacity of the dissolved organic matter (DOM) as being a possible explanation (He *et al*., 2019). Smaller water bodies have a higher surface area to volume ratio, so they tend to maintain a higher DOM concentration. Trout Bog Lake is also influenced by a dense Sphagnum mat surrounding the open water, which leaches DOM with high humic content (Maizel *et al*., 2017) that we expect to support the high electron accepting capacity. While some EET proteins in Trout Bog might be relics of ancient metabolisms, their abundance across genera instead suggests active use. The aforementioned observations and the scarcity of studies on EET in the water columns of lakes led to our suspicions that there may be more electroactive metabolism(s) in small water bodies than currently realized.

External electron acceptors like Fe(III), other metals, and complex forms of DOM are known substrates for heterotrophic EET as a terminal electron acceptor for respiration, in other words, electrogenesis or reductive EET (redEET) (Lovley *et al*., 1999; Lipson *et al*., 2010; Roden *et al*., 2010). All such acceptors might be used at some frequency planktonically in Trout Bog. EET can also be used to access external electron donors (Ross *et al*., 2011; Rowe *et al*., 2015), for example, as an electron source for reducing CO_2_ into biomass. Phototrophic isolates like some *Rhodopseudomonas palustris* strains are capable of electrotrophy or oxidative EET (oxiEET), accepting extracellular electrons from Fe(II) and electrodes, and some have been shown to be capable of growing by cryptically cycling photoreduced iron–organic matter complexes (Guzman *et al*., 2019; Peng *et al*., 2019). Cryptic cycling, defined as transformations unobserved due to quick turnover, likely occurs any time there is a regenerable redox-active substance, whether *via* photoreduction, across steep oxygen gradients in stratified systems, or *via* commensal metabolisms. EET may enable cryptic electron cycling directly and through otherwise inaccessible extracellular chemicals like some DOM. However, DOM-involved EET has remained largely unconsidered with a few notable exceptions (Berg *et al*., 2016; Lau *et al*., 2017; He *et al*., 2019; Li *et al*., 2020) in planktonic systems.

A metagenomic study showed that dominant planktonic phototrophs of the class Chlorobia in Trout Bog contain multiple copies the extracellular Fe(II)-oxidizing protein Cyc2 (He *et al*., 2019), much like many of its cultured and uncultured and globally dispersed relatives (He, Barco, *et al*., 2017; Garcia *et al*., 2021). However, it is difficult to determine what these uncultured Chlorobia are predominantly using for an electron donor because of their various oxidoreductases. They could potentially use sulfide, hydrogen, Fe(II), or reduced DOM via cryptic cycling of any or all of these substances. If, as it seems in Trout Bog, other lakes have phototrophic or chemotrophic electrotrophs living alongside electrogens and especially if DOM is a substrate or mediator for electrotrophs, then there may be an unconsidered regime of often-cryptic metabolism occurring in small lakes all across the globe.

Regardless of whether the electron exchange inferred from the high abundance of oxidative and reductive EET proteins in Trout Bog bacteria occurs directly with DOM or through DOM–metal cycling, questions remain: Are other similar bodies of water as inundated with EET-capable microbes, and is it related to DOM concentrations or quality? How might planktonic metabolisms such as anoxygenic photosynthesis, respiration, and others connect through EET? What environmental parameters lead to favorable conditions for planktonic EET metabolism—DOM quantity, iron, sulfur? To get a foothold on these questions, we performed a broad metagenomics analysis.

We hypothesized that DOM enriches for planktonic electroactive microorganisms in small boreal lakes by acting as a substrate and mediator for EET (Fig. 1). We therefore predicted that one would observe a significant positive correlation between a given lake’s DOC (dissolved organic carbon, a measure of DOM quantity) and the propensity of bacteria in that lake to house EET genes. To test this hypothesis, we correlated metagenomic and environmental characteristics of non-sediment, planktonic samples from 36 small boreal freshwater lakes, such as bog lakes and thermokarst lakes, in relation to EET genes and their prevalence across the inhabiting diversity of bacteria.

**Figure 1.**
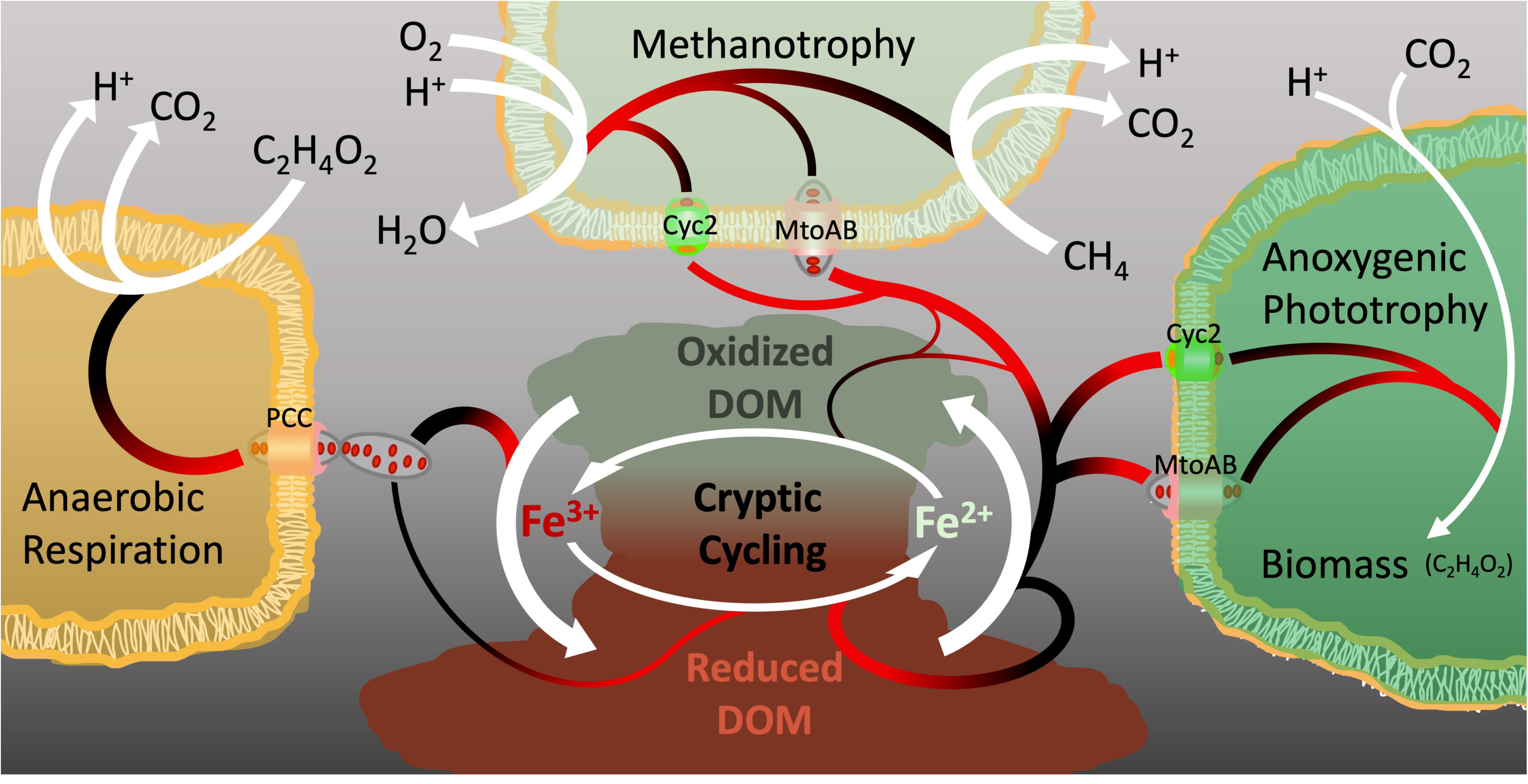
Conceptualized cryptic electron cycling and putative role in methane metabolism in small boreal lake water columns. Chemical transformations are shown with white arrows. Electrons flow from black to red.

## RESULTS

### Putative EET proteins were found in roughly half of all bacterial genomes analyzed from planktonic boreal lake samples

We compiled a dataset of 5569 Metagenome-Assembled Genomes (MAGs), each over 50% complete and less than 10% contaminated, from 190 metagenomic assemblies from the water columns of 36 small boreal lakes in order to compare the planktonic representation of EET genes in lakes of similar yet contrastable features. These included thermokarst ponds of various ages from three to 60 years since thaw, sheltered bog lakes characterized by *sphagnum* moss growth in a surrounding bog mat, and a range of other small boreal lakes (Supp. Tbl. 1). The original DNA samples were collected from various water column depths representing niches defined by light, temperature, and redox status. The MAGs were clustered by 95% ANI (average nucleotide identity) to form 2552 metagenomic Operational Taxonomic Units (mOTUs) (Buck *et al*., 2021). We then defined “OTUs” to be lake-specific (i.e. counting each mOTU once per lake), yielding 3030 OTUs.

We developed the computational workflow “Find EET” (FEET) to identify putative EET protein-encoding genes within genomes (hereafter called EET+ organisms, acknowledging that the actual EET capability of each organism must be verified experimentally). The workflow is based on the bioinformatics tool FeGenie (Garber *et al*., 2020) but adds an automated version of the outer-surface MHC and porin–cytochrome *c* protein complex (PCC)-identification approach described previously (He *et al*., 2019). Among the 3030 OTUs from the full dataset, 50% contained at least one protein found by FEET, and 40% had two or more. We note that a few of FeGenie’s Hidden Markov Model (HMM) (Baum and Petrie, 1966) hits should not be counted as promising EET evidence when found by themselves, including DFE variants and FoxABCYZ (Deng *et al*., 2018; Garber *et al*., 2020), which leaves 47% of OTUs with at least one protein that may enable EET. However, these numbers may be higher given more complete genomes.

The ability to transfer electrons to substances outside the cell is likely to co-occur with other strategies used for redox cycling. Thus, we asked which other inorganic electron donors and acceptors might be accessible to the EET+ organisms. The bioinformatic pipeline METABOLIC uses HMMs to make such predictions (Zhou *et al*., 2022) and showed that EET was not the only method for redox metabolism, in the form of non-EET oxidoreductases (Supp. Tbl. 3), genomically available to about 98% of our EET+ bacterial OTUs. However, EET was the only identified inorganic pathway for either oxidative or reductive redox metabolism for one or both sides of an OTU’s redox metabolism in 11% of these OTUs (Supp. Tbl. 6).

### The capacity for EET in water columns of small boreal lakes is significantly correlated to metabolic and environmental factors

All future mentions of correlation are positive unless stated otherwise. The variation in overall number and diversity of EET genes found in each lake made us curious about which organismal and environmental characteristics were associated with EET capacity. To quantify the propensity of a system to select for EET+ organisms, we calculated the ratio of EET+ OTUs relative to the total number of OTUs found in each lake. We also calculated the average number of EET proteins per OTU as a proxy for the propensity of a system to enrich for proficiency, redundancy, or flexibility in EET metabolism. By both proxies, oxiEET in water columns of boreal lakes was strongly and significantly correlated to DOM concentration (Fig. 2A). Methane concentration was slightly but significantly correlated to the average number of redEET proteins per OTU and to the ratio and number of oxiEET proteins. We observed several other expected and unexpected correlations (Fig. 2A, Supp. Fig. 5&6), as discussed below.

**Figure 2.**
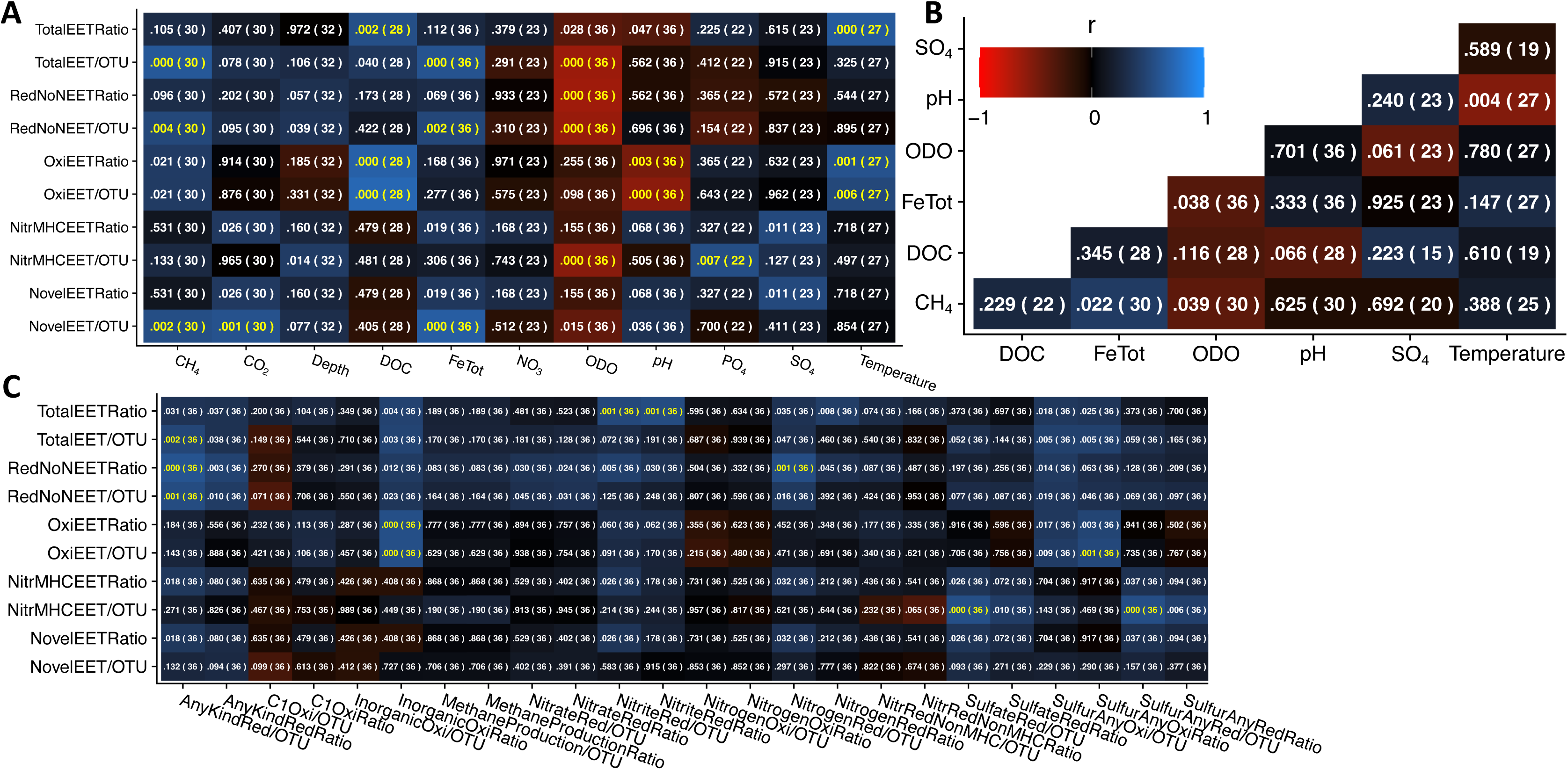
Correlations between environmental parameters (B) and EET values (A) and other oxidoreductase values (C) of 36 boreal lakes. Categories of protein values are delineated by “Ratio” or “/OTU” respectively standing for the ratio of OTUs with at least one of the given kind of protein or the average per OTU. Other values represent averages over available samples and data. For bog lakes involved in Long Term Ecological Research (lter.limnology.wisc.edu), recent (2018–2020) environmental data was subsampled by depth. Heatmap color represents Pearson correlation coefficient, r. Unadjusted p-values are out of “( N )” available corelates. Yellow text indicates a significance of p<.05 when adjusted by the Benjamini-Hochberg method. Statistical values and heatmaps were generated with the program R. Non-EET oxidoreductases were evaluated using METABOLIC (Zhou *et al*., 2022) whereas EET proteins were evaluated with the FEET pipeline. Category titles mean as follows: “TotalEET” = any putative EET protein; “RedNoNEET” = EET expected for reductive metabolism but excluding putative nitrate or nitrite reductase MHCs; “OxiEET” = EET expected to involve oxidative metabolism; “NitrMHCEET” = nitrate or nitrite reductases but only those that are identified as MHCs; “Novel” = outer membrane MHCs that were not classified as nor were 80% similar to any known EET protein; “C1Oxi” = single carbon compound oxidation; “NitrRedNonMHC” = Nitrate or Nitrite reductases that were not identified by FEET to be MHCs; “FeTot” = total iron and/or Fe(II) + Fe(III) measurements; “ODO” = optical dissolved oxygen “DOC” = dissolved organic carbon. Included HMMs and full category descriptions are listed in Supplementary Table 3.

### We observed putative EET proteins in most small boreal lake methylotrophs, especially methanotrophs, and these often coincided with carbon and nitrogen fixation proteins

Of the 121 Methylococcales OTUs (taxonomically expected methanotrophs) 83%, or 88% of those that were EET+, housed Cyc2 or another putative oxiEET protein. Of the 113 *Methylophilaceae* OTUs (expected methylotrophs) 58%, or 65% of EET+, had oxiEET. One possible usage of electrons is to support carbon and nitrogen fixation. Of the Methylococcales with oxiEET, 51% harbored nitrogen fixation marker proteins, and another 8% had carbon fixation markers, whereas only 35% of the few without oxiEET had N-fixation markers and none had C-fixation markers. Only one *Methylophilaceae* OTU had a N-fixation marker, and it did not have oxiEET. Of those with oxiEET, however, 12% did have C-fixation indicator proteins. Of the 16% of EET+ Methylococcales that had multiple redEET proteins as defined (Supp. Tbl. 4), these proteins were generally DFE variants or MtrB with MtoA; so, neither specifically indicates EET-mediated respiration (Hartshorne *et al*., 2009; Liu *et al*., 2012). The porin MtrB is defined as both redEET and oxiEET because, with our methods, it indistinguishable from the MtoB porin (Supp. Tbl. 4). Only two *Methylophilaceae* had methane monooxygenases, and both were EET+. Of Methylococcales with oxiEET proteins 82% had a methane monooxygenase, and of those without oxiEET, 60% had a methane monooxygenase (Supp. Tbl 6). We note that some MAGs within *Methylophilaceae* and Methylococcales were reported as EET+ (Tanaka *et al*., 2018; He *et al*., 2019; Tsuji *et al*., 2020; Yang *et al*., 2020 – respectively found: *Methylococcus capsulatus* redEET, *Methylotenera* oxiEET, *Methylomonadaceae* oxiEET, *Methylophilus* redEET).

### EET proteins in small boreal lakes were found in bacterial groups not yet documented to exhibit electroactivity

Historically familiar lake bacteria including *Polynucleobacter* and *Limnohabitans* (Newton *et al*., 2011) and several other taxa not previously considered to be capable of EET were represented by OTUs with proteins that strongly indicate EET metabolism (Fig. 3, Supp. Tbl. 2). Three Nanopelagicales (acI Actinobacteria) out of 358 Actinobacteria OTUs had between one and three EET proteins. As noted above, we were not the first to identify EET+ members of Methylococcales and *Methylophilaceae*, but we also identified additional EET+ taxa like *Methylomonas, Methylovulum, Methylopumilus,* and other *Methylococcaceae*. All the EET+ taxa are summarized in Supplementary Table 2 (Ehrenreich and Widdel, 1994; Suzuki *et al*., 2006; Croal *et al*., 2007; Qiu *et al*., 2008; Karthikeyan *et al*., 2012; Johnson *et al*., 2014; Li *et al*., 2015; Lino *et al*., 2015; Yu *et al*., 2015; He, Barco, *et al*., 2017; He, Stevens, *et al*., 2017; Eichorst *et al*., 2018; Jiang-Hao *et al*., 2019; Kjeldsen *et al*., 2019; Yang *et al*., 2020; Koskue *et al*., 2021), including more that have not been previously considered capable of EET.

**Figure 3.**
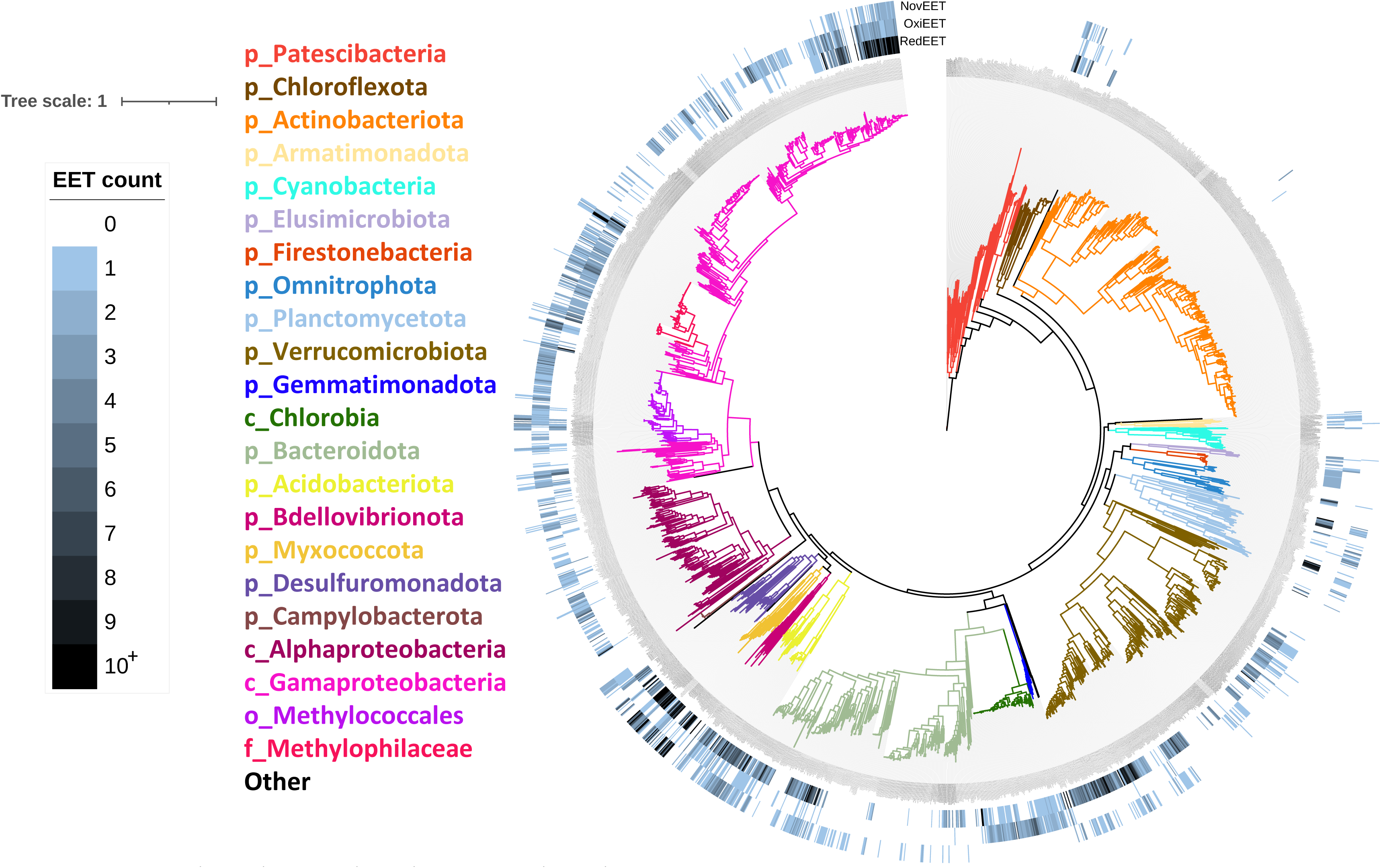
Putative novel (NovEET), oxidative (OxiEET), and reductive (RedEET) EET protein counts across phylogenetic tree of small boreal lake bacteria. OxiEET and redEET protein counts do not include MtrB and only include those specifically in the oxidative or reductive categories as designated (Supp. Tbl. 4). A 5040 amino acid-long sequence alignment of concatenated GTDB-tk markers genes found in the 2536 out of 2552 mOTUs which did not have identical marker sequences was used by RAxML-HPC BlackBox (Kozlov *et al*., 2019) on CIPRES (Miller *et al*., 2010) to generate a tree file set to use optimal bootstrapping. Tree visualization was performed using iTOL v5 (Letunic and Bork, 2021). GTDB-tk-assigned taxonomic level is designated by “p_” for Phylum-level and so on, and taxa colors are listed in clockwise order from the root.

### 606 novel putative EET Multiheme Cytochrome protein clusters were identified in taxa previously known and unknown to be electroactive (Fig. 3)

We refer hereafter to these EET MHC clusters as “novel clusters.” For each novel cluster (Supp. Tbl. 5) we have no evidence for homology with each other, MHC nitrate nor nitrite reductases, nor any known EET protein, using a cutoff of 20% amino acid sequence identity. This cutoff was chosen as a coarse balance between maintaining likely semblance in structure or function within clusters yet having a useful level of grouping for analysis. Taxa in which novel clusters were found include the following (numerically summarized in Supp. Tbl. 2, Fig. 3, Supp. Fig. 1).

Members of the phylum Myxococcota (Supp. Tbl. 2) comprised the top several taxa with the highest average number of novel clusters, closely followed by Geobacterales, Desulfocapsaceae, and then Geothrix. Accompanying the Myxococcota novel clusters was a mix other EET proteins like DFE variants, MtrC, ExtABC, CbcL, and Cyc2. Notably, EET has been experimentally demonstrated in the Myxococcota member Anaeromyxobacteraceae (Marshall *et al*., 2009).

Members of Planctomycetota including Phycisphaerae and especially Planctomycetes, the class that includes annamox bacteria which were very recently shown to be capable of EET (Shaw *et al*., 2020), were found containing one or several novel clusters and often with no other identifiable EET protein, though occasionally—and more so with Phycisphaerae—DFE variants or CbcL. However, none of the Planctomycetota were found to have characteristic annamox proteins, HzoA or HzsA.

The cosmopolitan and freshwater-specific Nanopelagicales contributed OTUs carrying novel clusters. The common freshwater genus *Limnohabitans* contributed OTUs with one novel cluster accompanied by either Cyc2 or MtoA and MtrB. The cosmopolitan anoxic phototroph Chlorobia contributed an OTU with a novel cluster and Cyc2. Numerous other instances of novel clusters distributed in taxa are summarized (Supp. Tbl. 5).

## DISCUSSION

### We have provided evidence that DOM mediates EET in small boreal lakes

We hypothesized that DOM serves as both an extracellular electron donor and acceptor in lakes with high DOM concentrations, not simply as an electron acceptor for redEET as proposed for humic substances acting as a substrate or electron shuttle in sediment and soil (Heitmann *et al*., 2007; Roden *et al*., 2010) and lakes (Lau *et al*., 2017). In support of this hypothesis, our prediction—that DOC will correlate to the propensity of bacteria to house EET proteins—was true, but not for all forms of EET. While DOM can be an optional receiver of electrons from many respiration-linked EET proteins traditionally associated with iron reduction (Lovley *et al*., 1999; Gralnick and Newman, 2007), DOM has not yet been demonstrated to be a direct donor for autotrophs which would be expected to require net electron input (*i.e.* obtained from extracellular donors) to reduce CO_2_ into biomass. Yet, oxiEET is significantly correlated with DOC while redEET is significantly correlated with iron, and not the reverse (Fig. 2A). Though these correlations may suggest DOM acts as a direct electron donor for oxiEET, as we propose, this inference cannot be made without acknowledging the correlations may also be due to DOM supporting oxiEET organisms in other ways. DOM could promote oxiEET by promoting the activity of redEET both as an electron-shuttling agent for dissimilatory Fe(III) reduction and as DOM is fermented into small organic acids that can be respired, fueling redEET and thereby regenerating Fe(II) for oxiEET. Also, cryptic photo-regeneration of Fe(II)-ligands as evidenced previously (Caiazza *et al*., 2007; Van Trump *et al*., 2013) likely adds to materials available for electrotrophs. As expected and reflected in its correlation to redEET, iron in high concentrations will likely (depending on pH) have an insoluble, oxidized fraction and will also increase Fe(II)-ligand-induced DOM photolysis (Caiazza *et al*., 2007), both of which likely enrich for redEET.

The ability of electrotrophs to use reduced forms of DOM as electron donors may reflect structural aspects of oxiEET proteins. For example, in the known oxiEET systems, an analogue for the redEET protein, MtrC—an extracellular protein which may span the negatively charged lipopolysaccharide layer (Edwards *et al*., 2020)—is lacking or not identified which may be due to the propensity of Fe(II) to be soluble only in this reduced form and therefore able to diffuse through the lipopolysaccharide layer. As Fe(II) is oxidized, precipitating Fe(III) can present physiological challenges for some bacteria through encrustation, though perhaps not as much at low pH like in many of these small boreal lakes as contemplated previously (Hegler *et al*., 2010; Bryce *et al*., 2018). In contrast, we expect DOM for the most part to be soluble in either reduced or oxidized forms. We do indeed observe a correlation between low pH and DOC (Fig. 2B), both of which are correlated to oxiEET (Fig. 2A). It is this combination of factors that may allow pelagic electrotrophs access to DOM and iron’s redox chemistry. These factors seem particularly important for photolithoautotrophs using oxiEET. When using DOM or Fe(II) as extracellular electron donors to fix CO_2_ into biomass (Fig. 1), they would also require a net assimilation of hydrogen, perhaps as protons aided by low pH, so we speculate. Indeed, in the dataset (Fig. 3, Supp. Fig. 3), other than expected methanotrophs as we later discuss, possible and probable phototrophs in Chlorobia, Alphaproteobacteria, and Gammaproteobacteria comprised much of the small lake OTUs with oxiEET proteins. OxiEET was also found within a variety of organoheterotrophs, which corresponds with previous research. The ubiquity of anaerobic respiring organisms in soils and sediments that are capable of coupling DOM use as an extracellular electron donor to the reduction of inorganics like nitrates has been previously noted (Coates *et al*., 2002; Roden *et al*., 2010), likely using oxiEET proteins.

### EET in the water column has implications for methane emissions

Only eight out of 5569 MAGs analyzed here were predicted to encode machinery for methanogenesis, among only 21 archaeal MAGs that passed the quality cutoffs. This, along with dissolved methane measurements in similar systems (Rissanen *et al*., 2021; van Grinsven *et al*., 2021), suggests that the majority of methanogenesis is occurring in sediments rather than the water column. The observed correlation between iron and methane, both of which are negatively correlated with dissolved oxygen (DO) (Fig 2B), is unsurprising because methanogenesis occurs typically when available iron and other electron acceptors are in the reduced state and Fe(II) is more soluble than Fe(III). Considering this and that iron correlates as it should to redEET, there may be noncausal correlation between redEET and methane. However, oxiEET appears to be independently correlated to methane concentrations (Fig. 2A&B, Supp. Fig. 5&6), and a simple explanation for this is that methanotrophs with oxiEET proteins are responsible.

However, the observed ubiquity of methylotrophs with oxiEET mechanisms is surprising considering methane or methyl groups are rather electron-rich, and an electron-intake mechanism like oxiEET would seem redundant unless many of these organisms are frequently not using single carbon compounds as the only electron source. One possibility is that the metabolism of these methanotrophs is more flexible than the classically understood methane oxidation model (Chistoserdova and Lidstrom, 2013). One might suspect that some require additional electron intake for carbon fixation or even nitrogen fixation. At least for Methylococcales with oxiEET, about 60% had nitrogen fixation genes or carbon fixation genes, but this explanation overlooks how the other 40% might use oxiEET (Supp. Tbl. 6). Considering the aforementioned ubiquity and that most Methylococcales did indeed have some form of methane or methyl monooxygenase, we suggest planktonic methanotrophic EET plays a role in regulating greenhouse gas emissions by supplementing metabolism when methane is limiting and therefore growing or sustaining their population until methane is not limiting. In particular, they may use oxiEET for energy generation by coupling to the reduction of nitrogen compounds, especially considering 82% of Methylococcales and 40% *Methylophilaceae* with oxiEET also have at least one reductase for nitrate, nitrite, or nitric oxide (Supp. Tbl. 6). Otherwise, they might couple oxiEET to oxygen reduction when operating in a microaerophilic zone.

Methylococcales and *Methylophilaceae* oxiEET genes were typically Cyc2 and/or the MHC MtoA. MtoA was nearly always accompanied by the outer membrane porin MtrB—which is not distinguished from MtoB in FEET (Garber *et al*., 2020), consistent with Methylococcales observed in Rissanen *et al* (2021). Because MtoA, though likely used for oxiEET, has been observed to operate in redEET (Liu *et al*., 2012), one might suspect they can couple redEET to methane oxidation like in Yang *et al* (2020). However, we observed no difference in the proportion of methylotrophs with Cyc2 or MtoA regardless of sample depth, including Methylococcales well below the oxycline, and if being used primarily for redEET coupled to methane oxidation, one may expect MtoA to be less common than Cyc2 in methylotrophs lower in the water column where there is more methane.

Furthermore, we found the strength of the correlation between temperature and oxiEET (Fig. 2A) surprising because temperature affects the rate of all metabolism, not just oxiEET. Perhaps this correlation is due to oxiEET’s role in autotrophy and its particular temperature dependencies (García-Carreras *et al*., 2018). Whatever the underlying mechanisms, this suggests that as thermokarst continue to warm(Zandt *et al*., 2020), oxiEET may become more prevalent. Furthermore, if this is due to methanotroph activity, EET may play an even larger role in controlling methane emissions.

Electrogenic respiring organisms compete with methanogens for small organic compounds such as acetate (Heitmann *et al*., 2007; Lau *et al*., 2017; Zakaria and Dhar, 2021), and their activity may be further enabled by mutualistic interactions with oxiEET by methanotrophs, chemotrophs, or autotrophs, acting as control on methane emissions independent of methane oxidation (Fig. 1). Examples of mutualistic behavior that may support ecologically competitive populations of respiring EET bacteria include the production of oxidized respirable materials or labile organic matter upon lysis or as autotrophic exudates. We expect these controls to be most relevant in upper stratified anoxic zones. Most known electrogens are anaerobes, partly due to oxygen binding heme groups in MHCs which makes them unable to donate electrons unless they have a compensating mechanism, and oxiEET mechanisms may be subjected to the same pressures. As expected, DO significantly correlated negatively to redEET genes. However, there was a lack of significance and an unexpected weakness in the negative correlation between DO and oxiEET genes. This is likely in part because many chemotrophic iron-oxidizers require oxygen as a TEA, but they are usually microaerophilic. We also considered that Chlorobia and other anoxygenic phototrophs, the other major boreal hosts of oxiEET, are often abundant just below the oxycline (Brand *et al*., 2016). When epilimnion sample collection integrates multiple depths or when there is mixing in the surface layers, either may cause organisms that live in low-oxygen environments to be found in the high-oxygen samples. Indeed, many methanotrophs in oxygenated samples had oxiEET proteins.

### Much like in sediments and soils, boreal lake EET is often connected to the sulfur and nitrogen redox cycles

Our results suggest that while EET is sometimes the only pathway available for redox metabolism, generally it is accompanied by non-EET oxidoreductases, as reflected in the strong positive correlations in Figure 2C. These include proteins enabling more commonly recognized energy generation modes such as nitrate respiration, sulfide oxidation, and ammonia oxidation. Notably, 84% of OTUs with identified respiratory pathways, EET or otherwise, had no identified inorganic oxidation pathway, implying that organotrophy dominates the microbial diversity of small lakes with high DOM. Still, similar couplings of EET to oxidoreductases have been observed in non-planktonic habitats (Osorio *et al*., 2013; Kawaichi *et al*., 2018). Building on such previous studies, we suggest that planktonic EET can cooccur or alternate with and/or couple to sulfur and nitrogen redox transformations.

As an example, oxiEET was more often found in small lakes where a higher proportion of organisms were capable of some manner of inorganic oxidation using donors such as ammonia or nitrite, and in particular sulfur, sulfide, sulfite, or thiosulfate—suggesting that bacteria may often alternate between or concurrently use oxiEET and such metabolisms. RedEET is more often found in lakes with a high proportion of organisms with proteins for nitrate/nitrite reduction, but also especially oxidation of sulfur forms (Fig. 2C)—suggesting redEET often cooccurs or alternates with nitrate/nitrite respiration yet, as observed previously in Lovely and Phillips (1994), couples to sulfur oxidation. The novel EET clusters (Supp. Tbl. 5) in particular may be involved with nitrite respiration, sulfate respiration, and/or iron metabolism, based on correlations to both oxidoreductases and environmental conditions conducive to these transformations (Fig. 2C).

Given the frequency of EET possibly being used as an additional redox source and the frequency of possible redox couplings to EET, we propose that EET may typically be a principal mode of microbial metabolism in small boreal lakes. However, to investigate precisely how often EET functions as a principal rather than as a backup metabolic mode in these systems, future studies based on transcriptome profiles, sensitive in situ electrochemical analyses, and/or pure cultures will be necessary.

### EET protein count reflects the expected ecology and metabolism of OTUs with implications for planktonic taxa in small boreal lakes

The distribution of EET proteins in genomes across the full diversity of MAGs in the dataset confirmed some expectations but also yielded some surprises (Fig. 3, Supp. Fig. 1–4). For example, OTUs from the phylum Cyanobacteria, other than one OTU in the class Vampirovibrionia, had no identifiable EET proteins which makes sense given their abundant electron donor, H_2_O. Regarded thus far as non-photosynthetic, Vampirovibrionia are not representative of the classically understood Cyanobacteria nor as well studied (Grettenberger *et al*., 2020). This raises questions about how or why Vampirovibrionia carry the oxiEET proteins Cyc2 and FoxY. The widely distributed Candidate Phylum Radiations (CPR), Paceibacteria and Saccharimonadia, were never found harboring EET proteins. While notable, this is not unexpected considering the observed tendency of CPR to lack even some of the most conserved bacterial components (Hug *et al*., 2016).

We also noted the lack of EET proteins in the class Actinobacteria. Of the 647 Actinobacteria MAGs, in only three did we find putative EET proteins. Considering that most Actinobacteria are organoheterotrophs relying on respiration for energy metabolism, we might have expected to find EET genes in this lineage simply because the capacity for EET is so markedly enriched across most genomes in our dataset. One possibility is that we may have had a search bias against finding EET in gram-positive bacteria like the class Actinobacteria because the study of gram-positive EET is relatively new with more to uncover mechanistically. At the time of analysis, the included EET proteins shown to confer EET in gram-positive bacteria were PplA, EetAB, FoxABC, and sulfocyanin (Garber *et al*., 2020). Supporting the possibility of this bias, in the same phylum Actinobacteriota but gram-negative class Thermoleophilia (Zarilla and Perry, 1986), eight out of 19 MAGs had EET proteins like CbcL homologues, DFE_0448, and outer surface MHCs.

Moderate numbers of oxiEET proteins seem associated with carbon assimilation while high copy numbers seem associated with energy generation. Organisms with oxidative EET protein counts two through six typically fell within Chlorobia, Alphaproteobacteria, and Gammaproteobacteria, most of which were expected to be methylotrophic or phototrophic based on their taxonomy (Fig. 3, Supp. Fig. 3, Supp. Tbl. 6). Some, particularly those with oxiEET count greater than six were actually expected to also respire (Chaudhuri and Lovley, 2003; Jangir *et al*., 2016; Gagen *et al*., 2019). However, a low number of oxiEET proteins, one to three, was associated with taxa expected to use a wide variety of metabolic strategies (Supp. Tbl. 2).

High numbers of redEET proteins seemed associated with energy generation through respiration as expected, though with some caveats (Fig 3, Supp. Fig. 4). For example, counts 1-6 represented a great variety of metabolisms, even including Chlorobia, a class expected to be obligately photoautotrophic. However, some evidence exists for substantial dark metabolism based on fermentation (Badalamenti *et al*., 2014) or respiration (Badalamenti *et al*., 2013). In Gammaproteobacteria, those with over 12 redEET proteins were in classes that include some photoheterotrophs. Some Phycisphaerae were found with high numbers of redEET, mostly DFE variants or CbcL, and they were rather rarely—only three out of 139 OTUs—found containing any putative oxiEET protein. One OTU in Bacteriovoracia, a prominent class whose known members are predatory, was found with 16 reductive EET proteins (Supp. Tbl. 4), four CbcL and 12 DFE variants, and one oxidative protein MtoA. Other OTUs in the similarly predatory class, Bdellovibrionia (Bratanis *et al*., 2020; Ezzedine *et al*., 2020), were found with several of both oxidative and reductive EET proteins. Thus, whether they respire oxidized cellular components or extract electrons, predatory bacteria may similarly gain virulence using EET like some mammalian pathogens (Light *et al*., 2018).

## CONCLUSION

In support of our hypothesis that DOM in aquatic systems can act as a substrate and mediator for EET-based metabolism, we discovered that Trout Bog is far from unique in its EET-enriched ecology (He *et al*., 2019); high proportions of putative electroactive bacteria and high numbers of EET proteins are present in small lakes across the globe. We also found that environmental propensity for EET correlates to DOC and other environmental and metagenomic values. Our findings have implications for small lake metabolism, green-house gas emissions, and our general understanding of bacterial ecology.

We have discovered potential roles for unconsidered proteins and added to our understanding of EET’s place in the environment by providing an overview of correlative relationships between EET and environmental conditions in small lakes, and by considering likely and possible resulting ecological interactions. Future modeling of methane emissions from thermokarst sediments, as permafrost continues to melt and form new ponds, should consider the roles of cryptic cycling and EET generally in the overlying water column. One likely role includes the mediation by planktonic methanotrophic bacteria where increased temperature, low pH, microbially derived DOM, and possibly pond age may increase the vibrance of the methanotrophic population beyond that expected from increased methane. Similar factors are also likely to increase the influence of bacteria using oxidative EET for anoxygenic photosynthesis. As reflected in our analysis, such influence likely extends from the role that DOM plays in enabling cryptic redox cycling, sustaining bacterial populations that respire or generate oxidized DOM that can then be (re-)respired, coupled to organotrophy and/or chemotrophy, thereby outcompeting methanogens in the water column for small organic fermentation products.

Though EET allows some bacteria to use solid-state surfaces, we show here that EET likely plays important roles even planktonically. This dataset analysis does not include large lakes which themselves often have areas of anoxia where EET may or may not be relevant, nor does it include representation of small lakes in tropical areas. There are also more recently discovered EET systems that are not fully represented in our analysis. Our findings set the stage for future studies to investigate these possibilities. Such studies may benefit from the use of cyclic voltammetry, *in situ* electrical measurements, and metatranscriptomics to investigate metabolic activity and to further define electroactive metabolism in lakes.

## EXPERIMENTAL PROCEDURES

### Sample Site Characteristics

The 36 small boreal lakes included in this dataset all have less surface area than 2.1 km^2^. Some are thermokarst melt pond ranging from old, middle-aged, and young (Peura *et al*., 2020). Some are acidic bog lakes with sphagnum moss growing over the edges. The rest are small lakes with a variety of surroundings and characteristics (Supp. Tbl. 1). The dataset includes lakes in Canada, Wisconsin and Alaska USA, Sweden, and Finland. All samples included in this study were from the water column and not from the sediment.

### Metagenomic Data Collection

MAGs (Supp. Tbl. 6) from some lakes, as denoted (Supp. Tbl. 1), were generated and made publicly available by collaborators (Buck *et al*., 2021), from which we used one metagenome per each of 186 samples. Other MAGs generated by the following methods were assembled from numerous samples from Long Term Ecological Research, and they comprise four of the metagenomes that are publicly available (https://osf.io/qkt9m/). Each was quality filtered with fastp 0.20.0 (Chen *et al*., 2018) and individually assembled with metaSPAdes v3.9.0 (Bankevich *et al*., 2012). Each metagenome was mapped to each individual assembly using BBmap from the BBTools v38.07 suite with a 95% sequence identity cutoff (Bushnell *et al*., 2017). Differential coverage from mapping to all samples was used to bin contigs into metagenome-assembled genomes using MetaBAT2 v2.12.1 (Kang *et al*., 2015). Bins were quality assessed with checkM v1.1.2 (Parks *et al*., 2015) and dereplicated with dRep v2.4.2 (Olm *et al*., 2017). All taxonomies were assigned by GTDB-tk v0.3.2 (Chaumeil *et al*., 2020).

### Finding EET Pipeline (FEET)

To find and summarize EET protein-encoding genes this pipeline combines a homology-based search tool, FeGenie (Garber *et al*., 2020) modified with additional HMMs (Supp. Tbl. 4), with the following more generalized approach that may allow for the identification of novel EET genes. MHCs participate in EET in at least two forms, 1) as extracellular or outer membrane MHCs to interact with extracellular substrates (such as OmcE, OmcS, and OmcZ), and 2) as a periplasmic MHC inserted into a porin to form a PCC (such as MtrABC and PioABC). FEET searches for both cases. FEET first uses python to find MHCs using the following regular expressions for heme binding motifs: [C.CH], [C..CH], [C…CH], and [C………..[!=C]..CH] (http://www.python.org). We set FEET’s adjustable parameters so that for FEET to call an MHC, the amino acid sequence must contain at least three [C..CH] motifs and at least five total aforementioned motifs. It then must be predicted by Cello V2.5 (http://cello.life.nctu.edu.tw/) likely to be an extracellular or outer membrane protein. Otherwise, it could be a periplasmic protein within eight open reading frames of a beta-barrel outer membrane porin (BB-OMP) predicted by a HMM or another extracellular or outer membrane-predicted MHC protein. The BB-OMPs in PCCs were also counted as EET proteins. All protein predictions were crosschecked between methods using FEET scripts. FEET pipeline performed here excludes a search for e-pili because at the time of analysis we lacked a way to distinguish PilA used as e-pili over Type IV secretion, however it has been since added to FeGenie, and future usage of FEET would also include it and any other recent additions. The pipeline and output is publicly available on Github (https://github.com/McMahonLab/FEET.git).

### FEET Performance

FEET pipeline performs as expected, finding EET proteins in model EET organisms and not finding EET proteins in model non-EET organisms. In order to ground-truth the FEET tool, we used it to analyze reference genomes of well-studied model organisms that are known or expected to contain EET proteins, including *Geobacter sulfurreducens* PCA – GenBank accession number GCA_000007985.2 and *Shewanella oneidensis* MR – GenBank accession number GCA_000146175.2. We also checked some genomes from organisms expected not to be capable of EET, including, *Chlorobaculum tepidum* TLS – GenBank accession number GCA_000006985.1, *Synechocystis sp.* PCC 6803 – GenBank accession number GCA_001318395.1. FEET found 54 EET proteins in *G. sulfurreducens* including one or more copies of CbcABL, OmcS, Cyc2, OmcFSZ, DFE_0461, DFE_0462, DFE_0463, DFE_0449, DFE_0448, ExtABCD, ImcH, Geobacter-associated PCCs, MtrA, MtoA, and 12 unidentified outer surface MHCs. FEET found 14 EET proteins in *S. oneidensis* including multiple copies of MtrABC, a DFE_0448, and a DFE_0465. *C. tepidum*, *Synechocystis sp*., and *E. coli* did not contain any identified EET proteins.

### Novel Sequence Curation and Crosschecking

Novel EET MHC amino acid sequences were clustered by 80 percent identity to infer homology using CD-hit (Li and Godzik, 2006) software with all the pipeline’s EET proteins as well as representative nitrate or nitrite reductase MHCs from the METABOLIC V3 pipeline (Zhou *et al*., 2022). Sequence identities were crosschecked with all other oxidoreductase HMM matches (Supp. Tbl. 3) from METABOLIC and the top NCBI BLAST (https://www.ncbi.nlm.nih.gov/) results (Supp. Tbl. 5). METABOLIC relies on unique HMMs as well as HMMs from TIGRFAM (Haft, 2003) and KEGG (Kanehisa, 2000) databases.

### Statistical Analysis

All Pearson correlation statistics were generated using R (R Core Team, 2014). Lake-wide statistical values for proteins are based on the presence/absence of OTUs and not normalized by read depth. Not normalizing to the read depth overrepresents uniquely binned populations as opposed to the dominating bacteria. Lake-wide averages and standard errors for environmental data represent all available samples’ accompanying metadata except for bog lakes involved in Long Term Ecological Research (lter.limnology.wisc.edu) recent (2018–2020) environmental data was subsampled by lake depth. For example, “Depth” (Supp. Tbl. 1, Fig 2A) represents the average sample depth for most lakes. Only MAGs over 50% complete and less than 10% contaminated as evaluated by CheckM (Parks *et al*., 2015) were considered, and the most complete and least contaminated MAG from a given lake was selected to represent the OTU in that lake. Except in Supplementary Figures 1–4, taxa bar sizes were based on MAGs and normalized to the number of samples taken from each MAG’s lake. This was to circumvent protein count discrepancies between MAGs in a given lake belonging to the same mOTU. Decisions to remove certain potentially available bog, thermokarst, or other lakes’ metagenomic datasets from the analyses were based on either not having the required DOC data, being a lake of surface area over 2.5km^2^, or, in the case of one Canadian thermokarst pond labeled “F5,” because of especially limited metagenomic data, retaining 36 bodies of water.

### Graphical Analysis

Heatmaps and bar charts were generated using R’s ggplot2 package (Wickham, 2009). The phylogenetic tree was generated using the 5040 amino acid-long sequence alignment output from GTDB-tk of concatenated GTDB-tk markers genes found in the 2536 out of 2552 mOTUs which did not have identical marker sequences. This alignment was used to generate tree files with RAxML-HPC BlackBox (Kozlov *et al*., 2019) on CIPRES (Miller *et al*., 2010) using the default optimal bootstrapping. Tree file and EET metadata visualization was performed using iTOL v5 (Letunic and Bork, 2021).

## SUPPLEMENTARY FIGURE LEGENDS

**Supplementary Figure 1.** Taxonomic proportions (A) and number (B) of OTUs with specific counts of novel putative EET proteins in small boreal lakes. These include MHC that genes do not cluster by CD-hit within 80 percent identity of any known EET HMM match in dataset of 36 bog and thermokarst lakes. Counts include some unidentified porins near outer surface MHCs. Taxonomies were assigned by the GTDB-tk database. Figures were generated in R using ggplot2. Column “X” includes any count including zero, representing the dataset’s taxonomy. Proportions less than 0.005 are not labeled. Some lower taxa were separated out from class level. Taxa with numerical GTDB-tk designations were combined at a higher taxonomic level. Full results are in Supplementary Table 6.

**Supplementary Figure 2.** Taxonomic proportions (A) and number (B) of OTUs with specific counts of all EET proteins in small boreal lakes. Taxonomies were assigned by the GTDB-tk database. Figures were generated in R using ggplot2. Column “X” includes any count including zero, representing the dataset’s taxonomy. Proportions less than 0.005 are not labeled. Some lower taxa were separated out from class level. Taxa with numerical GTDB-tk designations were combined at a higher taxonomic level.

**Supplementary Figure 3.** Taxonomic proportions (A) and number (B) of OTUs with specific counts of oxidative EET proteins in small boreal lakes. Taxonomies were assigned by the GTDB-tk database. Figures were generated in R using ggplot2. Column “X” includes any count including zero, representing the dataset’s taxonomy. Proportions less than 0.005 are not labeled. Some lower taxa were separated out from class level. Taxa with numerical GTDB-tk designations were combined at a higher taxonomic level.

**Supplementary Figure 4.** Taxonomic proportions (A) and number (B) of OTUs with specific counts of reductive EET proteins in small boreal lakes. Taxonomies were assigned by the GTDB-tk database. Figures were generated in R using ggplot2. Column “X” includes any count including zero, representing the dataset’s taxonomy. Proportions less than 0.005 are not labeled. Some lower taxa were separated out from class level. Taxa with numerical GTDB-tk designations were combined at a higher taxonomic level.

**Supplementary Figure 5.** Correlations between and EET values from 36 boreal lakes. Categories of protein values are delineated by “Ratio” or “/OTU” respectively standing for the ratio of OTUs with at least one of the given kind of protein or the average per OTU. Heatmap color represents Pearson correlation coefficient, r. Unadjusted p-values are out of “( N )” available corelates. Yellow text indicates a significance of p<.05 when adjusted by the Benjamini-Hochberg method. Statistical values and heatmaps were generated with the program R. Full EET protein category descriptions are listed in Supplementary Table 3.

**Supplementary Figure 6.** Correlations between environmental parameters and non-EET oxidoreductase values of 36 boreal lakes. Categories of protein values are delineated by “Ratio” or “/OTU” respectively standing for the ratio of OTUs with at least one of the given kind of protein or the average per OTU. Environmental values represent averages over available samples and data. For bog lakes involved in Long Term Ecological Research (lter.limnology.wisc.edu), recent (2018–2020) environmental data was subsampled by depth. Heatmap color represents Pearson correlation coefficient, r. Unadjusted p-values are out of “( N )” available corelates. Yellow text indicates a significance of p<.05 when adjusted by the Benjamini-Hochberg method. Statistical values and heatmaps were generated with the program R. Non-EET oxidoreductases were evaluated using METABOLIC (Zhou *et al*., 2022). Included HMMs and full category descriptions are listed in Supplementary Table 3.

## SUPPLEMENTARY TABLE LEGENDS

**Supplementary Table 1.** Lake-wide information. This table includes lake characteristics, lake-wide EET protein values and METABOLIC protein values and standard deviations when available.

**Supplementary Table 2.** Unique taxa characteristics and references. This table includes average EET protein values for each unique taxa. It also contains our designation of the quality of evidence that each unique taxa contains members suspected capable of EET. These designations occur under the header “EET Evidence Quality” with following category keywords “obviously” (some members contain several promising EET genes as defined in Supplementary Table 3), “likely” (highest suspect members contain one or two promising EET genes), “possibly” (highest suspect members contain two or three DFE variants or FoxABCYZ proteins but no other more promising EET genes), and “unknown” (contains just one of such “possible” EET genes). This table also contains whether each unique taxa has been found in a previous study to have or likely have EET capabilities under the header “Previous EET evidence” with the following keyword meanings: “yes” (we found studies that show EET in organisms in the genus or species), “no” (we did not find studies including genus or species doing EET; or taxonomic level is not defined enough to know), and “related”(organisms in the same order or family but not the particular genus or species were found in a previous study with EET).

**Supplementary Table 3.** Abbreviations and descriptions. In addition to abbreviations, this table lists the names of proteins for which HMMs were used in METABOLIC for each category.

**Supplementary Table 4.** Bit scores for FeGenie Hidden Markov Models and observed redox association. This table also includes additional conditionals for whether each HMM was considered as promising evidence of EET capability.

**Supplementary Table 5.** Novel cluster information. This table includes each unique novel cluster sequence, its percent identity to the reference sequence, and its number of heme motifs. Also included is which open reading frame the sequence represents, and what taxa it was found in.

**Supplementary Table 6.** Information unique to each metagenome assembled genome (MAG). This includes to what lake, sample, and mOTU each MAG belongs, its GTDB-tk taxonomy, genetic characteristics, and FEET and METBOLIC proteins.

## ACKNOWLEDGEMENTS

We give special thanks to Moritz Buck, Sarahi L. Garcia, Leyden Fernandez, Gaëtan Martin, Gustavo A. Martinez-Rodriguez, Jatta Saarenheimo, Jakob Zopfi, Stefan Bertilsson, and Sari Peura for early access to and setup with their dataset (Buck *et al*., 2021) to support this “pandemic project.” We thank the University of Wisconsin (UW)-Trout Lake Station, the UW Center for Limnology, and the John and Patricia Lane Award program for their invaluable support. We thank the U.S. National Science Foundation North Temperate Lakes Long-Term Ecological Research site (NTL-LTER DEB-1440297; DEB-0702395) for providing funding of the Microbial Observatory for long-term sampling of Lake Mendota and Trout Bog. We thank the U.S. Department of Energy Joint Genome Institute for sequencing and assembly (CSPs 394 and 2796). This research was also performed in part using the Wisconsin Energy Institute computing cluster, which is supported by the Great Lakes Bioenergy Research Center as part of the U.S. Department of Energy Office of Science. We are also thankful for fellowships provided through the department of Bacteriology at UW-Madison. We declare no conflicts of interest.

