## Supplementary Figure 6 for "Environmental predictors of electroactive bacterioplankton in small boreal lakes"

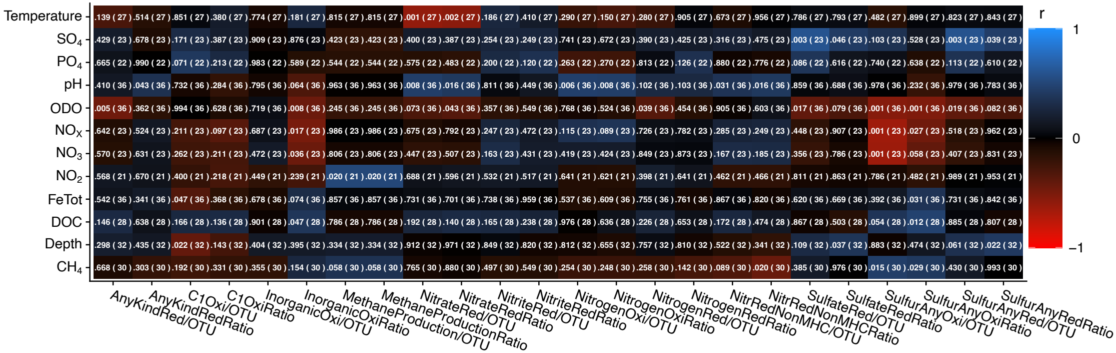


**Supplementary Figure 6.** Correlations between environmental parameters and non-EET oxidoreductase values of 36 boreal lakes. Categories of protein values are delineated by “Ratio” or “/OTU” respectively standing for the ratio of OTUs with at least one of the given kind of protein or the average per OTU. Environmental values represent averages over available samples and data. For bog lakes involved in Long Term Ecological Research (lter.limnology.wisc.edu), recent (2018–2020) environmental data was subsampled by depth. Heatmap color represents Pearson correlation coefficient, r. Unadjusted p-values are out of “( N )” available corelates. Yellow text indicates a significance of p<.05 when adjusted by the Benjamini-Hochberg method. Statistical values and heatmaps were generated with the program R. Non-EET oxidoreductases were evaluated using METABOLIC (Zhou *et al.*, 2022). Included HMMs and full category descriptions are listed in Supplementary Table 3.
