## Supplementary Figure 5 for "Environmental predictors of electroactive bacterioplankton in small boreal lakes"

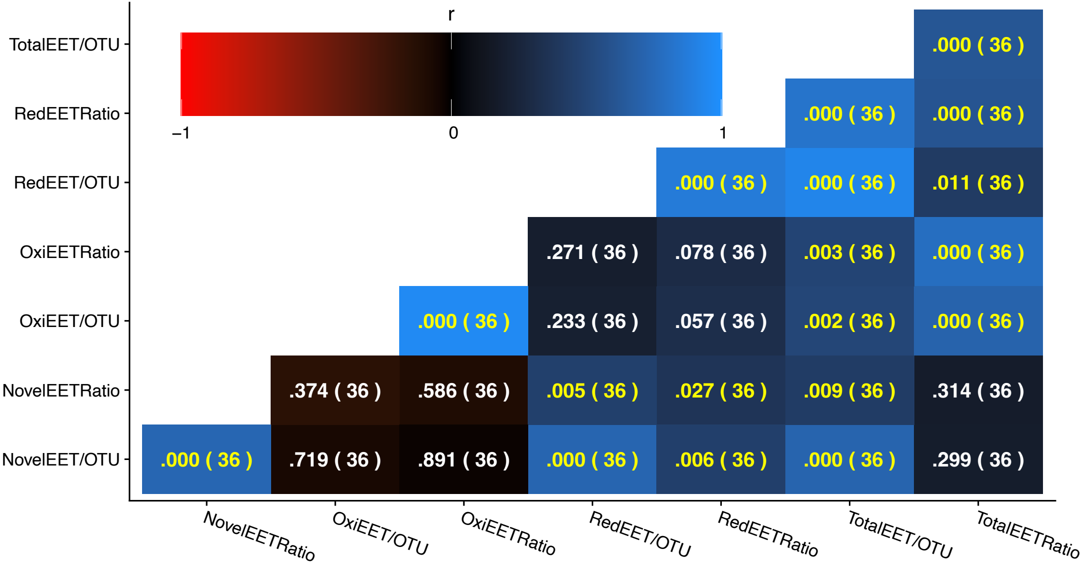


**Supplementary Figure 5.** Correlations between and EET values from 36 boreal lakes. Categories of protein values are delineated by "Ratio" or "/OTU" respectively standing for the ratio of OTUs with at least one of the given kind of protein or the average per OTU. Heatmap color represents Pearson correlation coefficient, r. Unadjusted p-values are out of "( N )" available corelates. Yellow text indicates a significance of p<.05 when adjusted by the Benjamini-Hochberg method. Statistical values and heatmaps were generated with the program R. Full EET protein category descriptions are listed in Supplementary Table 3.
