## Supplementary Figure 3 for "Environmental predictors of electroactive bacterioplankton in small boreal lakes"

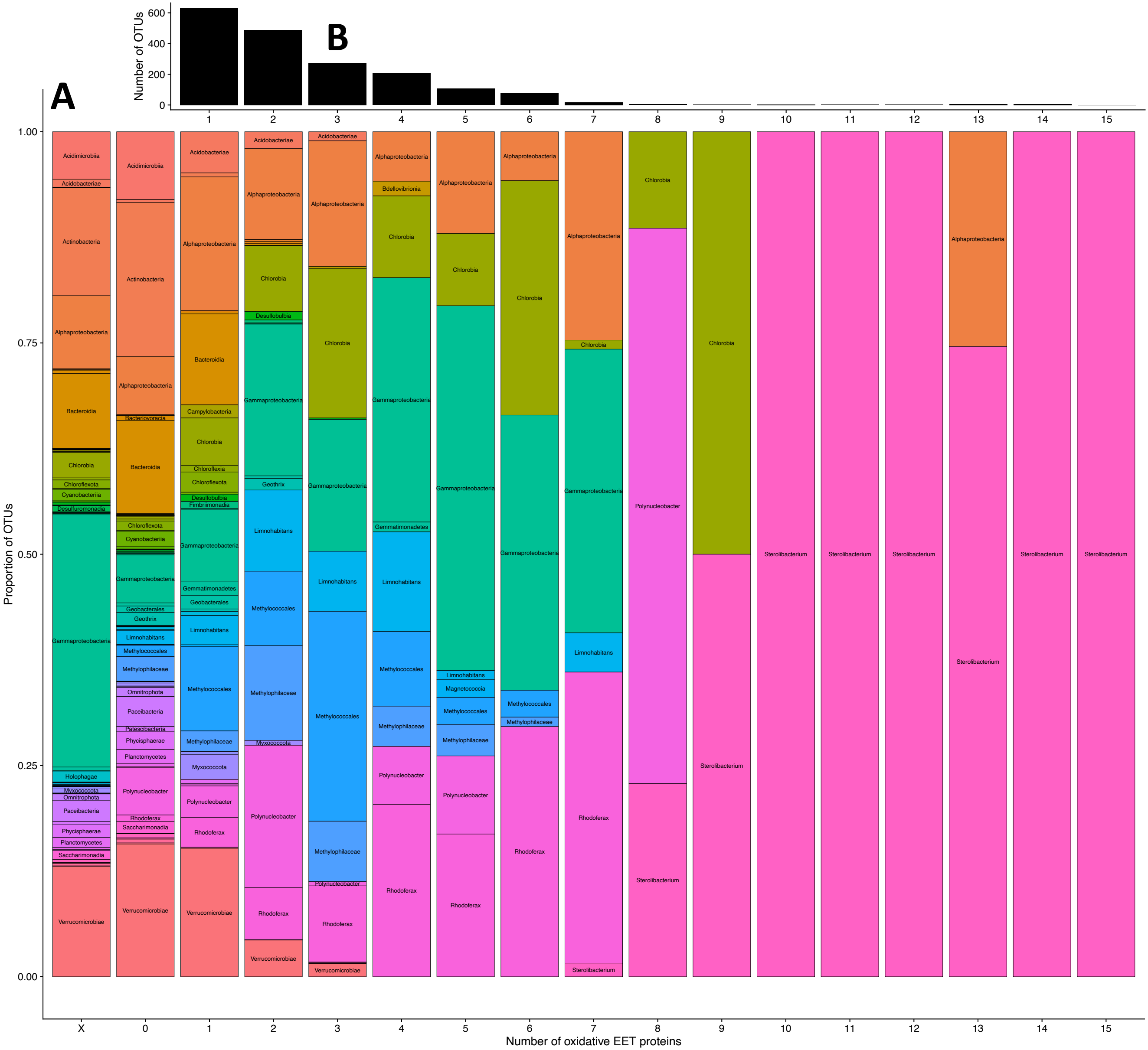


**Supplementary Figure 3.** Taxonomic proportions (A) and number (B) of OTUs with specific counts of oxidative EET proteins in small boreal lakes. Taxonomies were assigned by the GTDB-tk database. Figures were generated in R using ggplot2. Column "X" includes any count including zero, representing the dataset's taxonomy. Proportions less than 0.005 are not labeled. Some lower taxa were separated out from class level. Taxa with numerical GTDB-tk designations were combined at a higher taxonomic level.
