## Supplementary Figure 1 for "Environmental predictors of electroactive bacterioplankton in small boreal lakes"

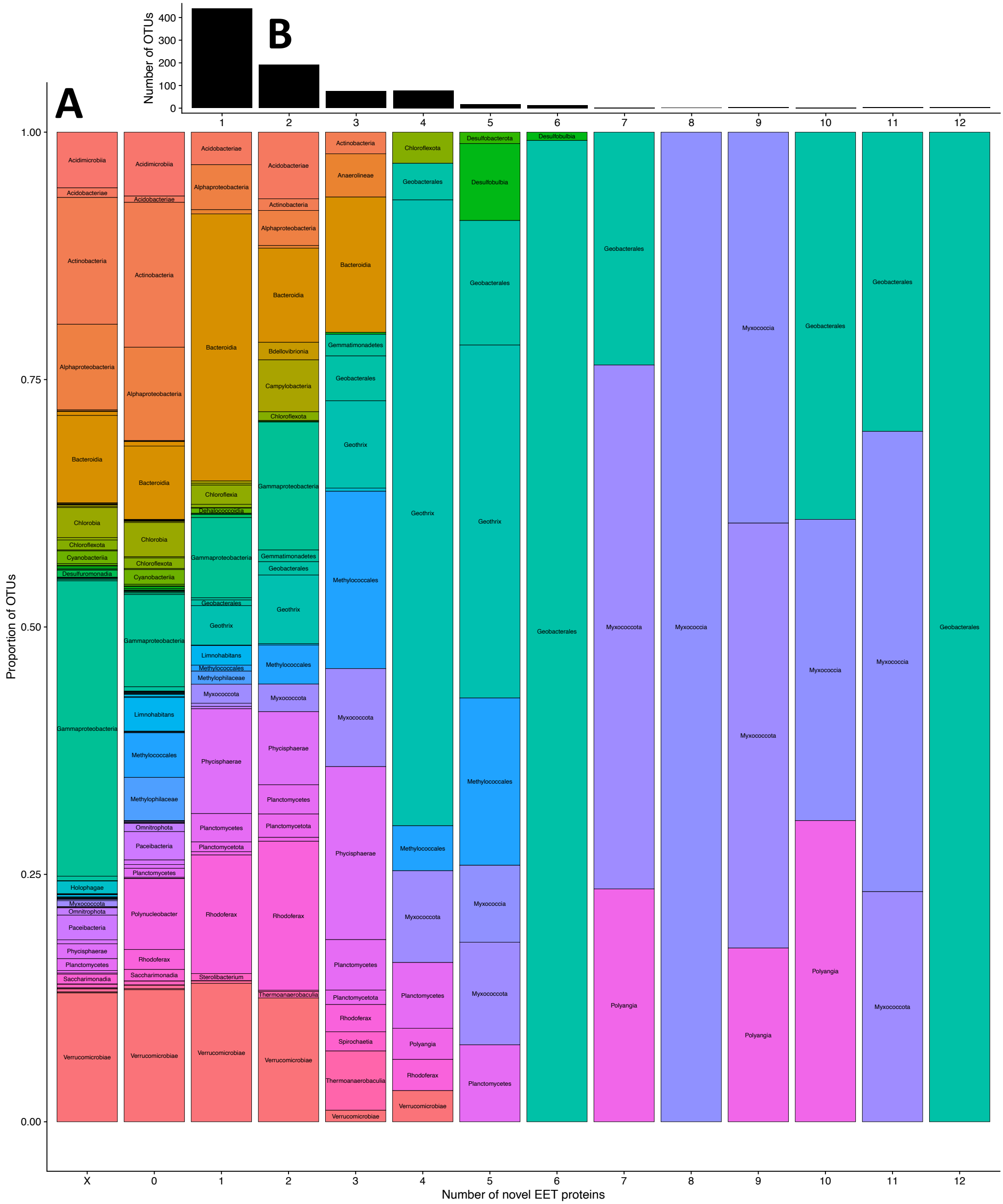


**Supplementary Figure 1.** Taxonomic proportions (A) and number (B) of OTUs with specific counts of novel putative EET proteins in small boreal lakes. These include MHC that genes do not cluster by CD-hit within 80 percent identity of any known EET HMM match in dataset of 36 bog and thermokarst lakes. Counts include some unidentified porins near outer surface MHCs. Taxonomies were assigned by the GTDB-tk database. Figures were generated in R using ggplot2. Column “X” includes any count including zero, representing the dataset’s taxonomy. Proportions less than 0.005 are not labeled. Some lower taxa were separated out from class level. Taxa with numerical GTDB-tk designations were combined at a higher taxonomic level. Full results are in Supplementary Table 6.
